# Identification and comparison of genes differentially regulated by transcription factors and miRNAs

**DOI:** 10.1101/803643

**Authors:** Amir Asiaee, Zachary B Abrams, Samantha Nakayiza, Deepa Sampath, Kevin R. Coombes

**Author notes:** These authors contributed equally (Amir Asiaee, Zachary B Abrams).

## Abstract

Transcription factors and microRNAs (miRNA) both play a critical role in gene regulation and in the development of many diseases such as cancer. Understanding how transcription factors and miRNAs influence gene expression is thus important to understand, but complicated due to the large and interconnected nature of the human genome. To help better understand what genes are being regulated by transcription factors and/or miRNAs we looked at it over 8000 patient samples from 32 different cancer types collected from The Cancer Genome Atlas (TCGA). We started by clustering the transcription factors and miRNAs using Thresher to reduce the number of features. Using both the mRNA and miRNA sequencing data we constructed linear models to calculate the coefficient of determination (R^2^) for each mRNA based on the Thresher cluster expression. We generated three types of linear models: transcription factor, miRNA and transcription factor plus miRNA. We then determined genes that were highly explained or poorly explained by each of the three models based on the genes R^2^ value. We performed downstream gene enrichment analysis using ToppGene on the sets of well and poorly explained genes. This identified differences in gene regulation between transcription factors and miRNAs and showed what groups of gene are differentially regulated.

## Introduction

Several different regulatory factors impact gene expression including transcription factors, methylation patterns, and miRNAs[1]. That these types of regulatory factors play a crucial role in gene expression and thus have an active role in disease progression has been well documented[2]. Thus, it is vitally important that we further our understanding of how genes are regulated in order to help treat why did variety of disease conditions. However, to date there are limited studies that have (1) identified the different types of genes that are regulated or expressed by these elements and (2) compared the differential effect of these regulatory elements on those genes[3]. In this article we present experimental results that focus on the comparative role of two particular regulatory elements: transcription factors and miRNAs. We selected these regulatory elements for two primary reasons. First, both elements regulate the abundance of messenger RNA in the cell[4]. Transcription factors do this by promoting transcription of a particular gene[5]. miRNAs regulate messenger RNAs by targeted binding and subsequent ubiquitination to degrade and destroy the messenger RNA[6]. Although miRNAs and transcription factors work differently, they both operate at the same relative stage of gene regulation[5]. Second, there is an abundance of publicly available data involving both mRNA and miRNA sequencing on the same patient cohort; most notably The Cancer Genome Atlas (TCGA)[7]. This enabled us to conduct a comparative study to evaluate the combined role of both transcription factors and miRNAs in regulating mRNA expression.

## Methods

### Data

As detailed previously, we analyzed data collected from TCGA using FireBrowse[8]. We identified 486 transcription factors as listed in the Transcription Factor Catalog[9], in 10,446 samples from studies of 33 different kinds of cancer in TCGA. We normalized the data using reads per kilobase per million (RPKM) and then performed a log2 transformation[10]. We also identified 8,895 patients from 32 cancer types that had miRNA sequencing data in TCGA using FireBrowse. We normalized the data using reads per kilobase per million (RPKM) and then performed a log2 transformation. We performed an additional filtering step by removing miRNAs that had a read count of zero in 75% or more of the patient population. This left us with 470 miRNAs across the 8,895 patients. We then identified the overlap between the transcription factor and miRNA cohorts. This left us with 8,895 patients who had both transcription factor and miRNA data along with mRNA sequencing.

### Software

The first step in our analysis was to identify clusters within both the miRNA and transcription factor data sets. This was done using the Thresher R package[11]. This R package generates one-dimensional clusters in principal component space. We used the MultiLinearModel function of version 3.1.6 of the ClassComparison R package to run our linear models[12]. We then calculated the coefficient of determination (R^2^) for each mRNA for each of the linear predictive models to gauge the prediction performance. The R^2^ the value is on a 0 to 1 scale and represents the proportion of variability explained buy the particular input factors in a model. Thus an R^2^ value of 0.4 means that 40% of the variability is explained by the model. The R^2^ values can determine both how influenced individual mRNAs our by regulatory elements and what type of regulatory elements, transcription factors and miRNAs, are having the greatest effect.

To perform our downstream gene enrichment analysis, we utilized the ToppGene website[13]. This web service allows users to input a set of genes and perform a large set of Fisher exact tests across a variety of curated biological pathways. The system can then tell a user which pathways are statistically significantly enriched for a given input set of genes.

## Results

### R squared distributions

We ran the Thresher algorithm over the set of miRNAs and the set of transcription factors independently. This produced 30 transcription factor clusters and 21 miRNA clusters. These clusters were then examined within a set of predictive linear models. We performed three sets of linear models: (1) transcription factor only, (2) miRNA only and (3) transcription factor plus miRNA. Each of these three models was run over the same data set of 8,895 patients and 20,289 mRNAs. Thus, each of our three sets of linear models was run for each of the 20,289 mRNAs so that an R^2^ value could be calculated for each mRNA. The distributions of R^2^ values for each model are shown in Figure 1.

**Figure 1.**
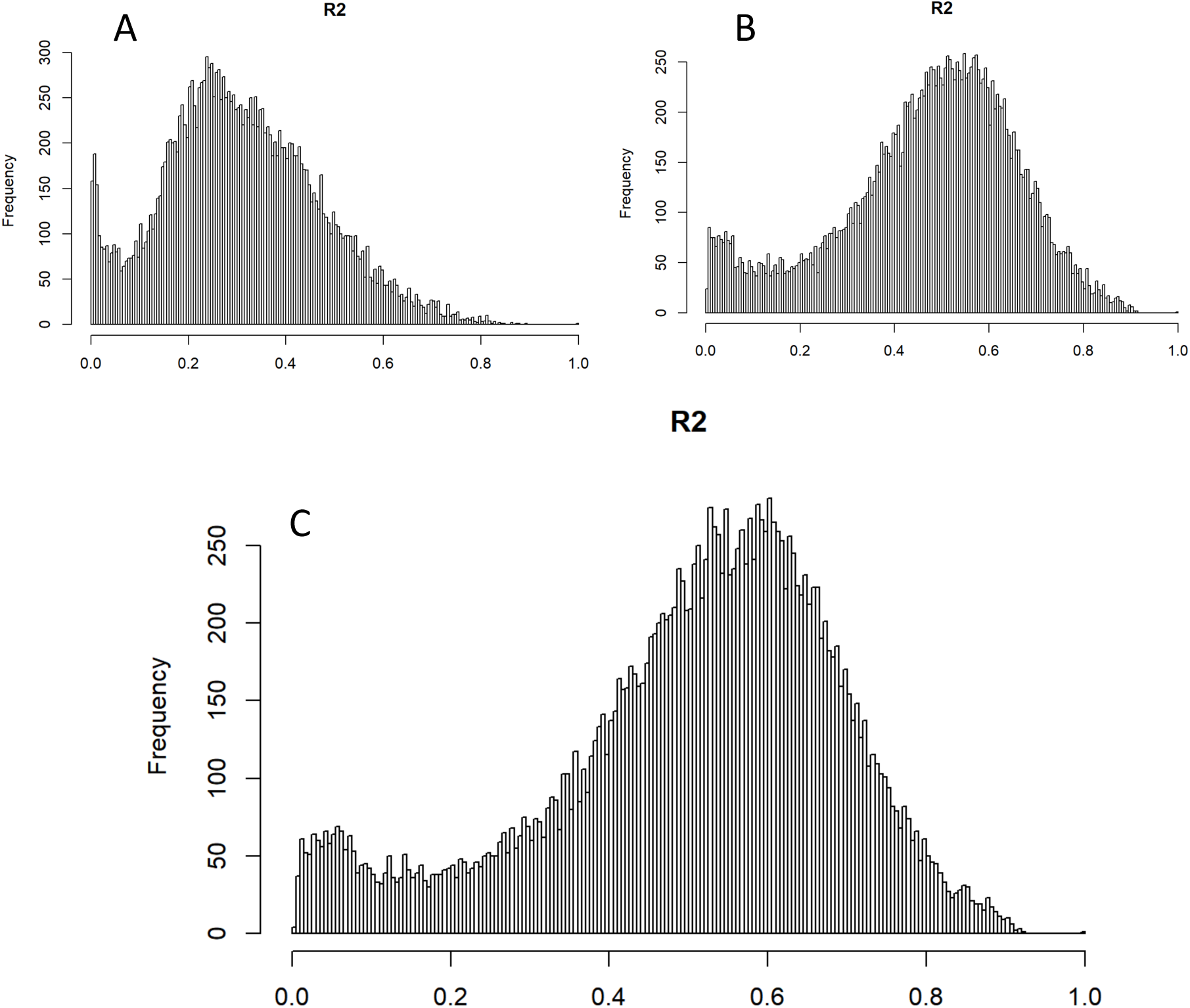
Distribution of R^2^ values across linear models. Panel A shows the miRNA model (mean = 0.30), B shows the transcription factor model (mean = 0.47) and C shows the joint model (mean = 0.50).

### ToppGene Enrichment

For each of the three models we identified the genes that had an R^2^ value of greater than 0.80 and less than 0.02. This identified genes that were either well or poorly explained by the models. These genes were then run through the ToppGene web service to identify gene pathways that were significantly enriched for each individual gene set. A subset of these results is shown in Table 1 (well explained genes) and in Table 2 (poorly explained genes). The complete lists are shown in Supplemental Table 1 and Table 2.

**Table 1.**
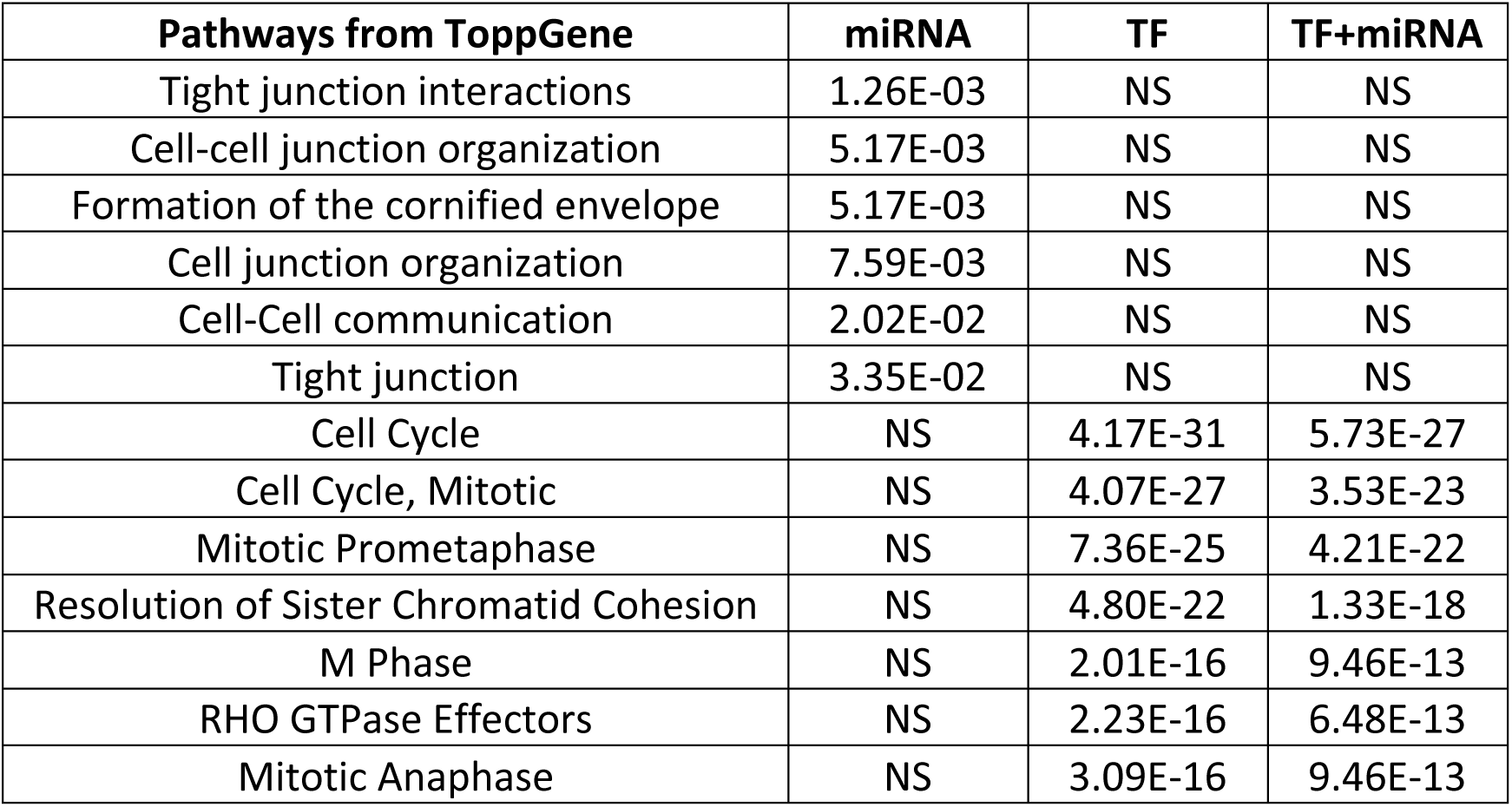
Gene enrichment for well explained genes (R^2^ > 0.80). p-values are calculated using a false discovery rate (FDR) correction Benjamini–Hochberg procedure. NS indicates non-significant. The main pattern involving genes that are well explained by miRNA expression is in cell-to-cell communication, such as the formation of junctions. This is in contrast to the transcription factor and joint model, which are dominated by cell cycle-related processes such as M phase and sister chromatid cohesion. These findings demonstrate that the classes of genes that are highly regulated by miRNAs and transcription factors do not have a high degree of overlap.

| Pathways from ToppGene | miRNA | TF | TF+miRNA |
| --- | --- | --- | --- |
| Tight junction interactions | 1.26E-03 | NS | NS |
| Cell-cell junction organization | 5.17E-03 | NS | NS |
| Formation of the cornified envelope | 5.17E-03 | NS | NS |
| Cell junction organization | 7.59E-03 | NS | NS |
| Cell-Cell communication | 2.02E-02 | NS | NS |
| Tight junction | 3.35E-02 | NS | NS |
| Cell Cycle | NS | 4.17E-31 | 5.73E-27 |
| Cell Cycle, Mitotic | NS | 4.07E-27 | 3.53E-23 |
| Mitotic Prometaphase | NS | 7.36E-25 | 4.21E-22 |
| Resolution of Sister Chromatid Cohesion | NS | 4.80E-22 | 1.33E-18 |
| M Phase | NS | 2.01E-16 | 9.46E-13 |
| RHO GTPase Effectors | NS | 2.23E-16 | 6.48E-13 |
| Mitotic Anaphase | NS | 3.09E-16 | 9.46E-13 |

**Table 2.** Gene enrichment for poorly explained genes (R^2^ < 0.02). p-values are calculated using a false discovery rate (FDR) correction Benjamini–Hochberg procedure. These results indicate that genes that are poorly explained by miRNA expression are also poorly explained using transcription factors. This may indicate that a different process, such as methylation, regulates these genes. Many of these pathways are associated with basic developmental biology such as the G protein-coupled receptor (GPCR) pathway.

| Pathways from ToppGene | miRNA | TF | TF+miRNA |
| --- | --- | --- | --- |
| Olfactory transduction | 6.77E-229 | 5.70E-58 | 2.11E-17 |
| Olfactory Signaling Pathway | 7.18E-223 | 3.92E-57 | 3.46E-17 |
| GPCR downstream signaling | 1.66E-154 | 3.56E-39 | 1.45E-11 |
| Signaling by GPCR | 2.75E-128 | 4.66E-31 | 8.27E-09 |
| Keratinization | 3.67E-43 | 3.29E-43 | 2.55E-37 |
| Defensins | 7.94E-16 | 5.99E-14 | 1.43E-11 |
| Developmental Biology | 1.15E-06 | 2.10E-13 | 4.31E-15 |

## Discussion

### Transcription factor and miRNA R^2^ values

There are some significant observations made by comparing the R^2^ values of the three linear models. First, the overall performance of the transcription factor only model (mean = 0.47) was greater than that of the miRNA only model (mean = 0.30). Second, adding the miRNA to the transcription factors did not significantly increase the model’s performance compared to the transcription factor only model. This may be because there are fewer genes (n = 32) that are well explained by the miRNA only model compared to the number of genes (n = 330) explained by transcription factors. Although the joint model had a mean expression of 0.50, it is likely that other regulatory elements not tested in this experiment contributed causally to the missing predictive strength. Events such as methylation are significant factors in gene expression and incorporating methylation data into this joint model in future research may improve the overall predictive strength of the model.

### Gene enrichment analysis

Large differences were evident in the gene enrichment results involving well explained genes between miRNA only and transcription factor only (Table 1). The joint model tracked strongly with the transcription factor only model. The miRNA model showed the genes that are highly regulated by miRNAs are involved in cell communication and junction formation. These results mirror other research that has linked miRNA regulation to tight junction proteins[14,15]. By contrast, the genes that are highly regulated by transcription factors are significantly involved in the cell cycle and other cellular reproduction/division systems. There is evidence in the literature that transcription factors play the key role in regulating cell division[16].

In contrast to the well explained genes, the set of genes that were poorly explained by the miRNA model and the transcription factor model are nearly identical (Table 2). Overall, genes that were poorly explained by all models are involved in basic developmental biology or olfactory receptor pathways. Olfactory receptors are an interesting finding, although it is known that olfactory receptors often come up as significant in a lot of gene enrichment studies due to the large number of different olfactory receptor genes[17]. However, the pathways associated with developmental biology and G protein-coupled receptors (GPCR) indicated that there are a large set of genes involved in fundamental biological processes that are virtually uncontrolled by transcription factors or miRNA regulation. This may be because these pathways often start signal cascades that may be influencing transcription factors to turn on and regulate other genes[18,19]. In essence the genes that are not controlled by miRNA or transcription factors may be influencing transcription factors and miRNAs, or be controlled by other gene regulatory elements like methylation. This finding illustrates the complexity of gene expression and the importance of continued research in the future.

## Supporting information

ToppGene raw results

